# Nanoscopic compartmentalization of membrane protein motion at the axon initial segment

**DOI:** 10.1101/046375

**Authors:** David Albrecht, Christian M. Winterflood, Thomas Tschager, Helge Ewers

**Affiliations:** Institut für Biochemie, Freie Universität Berlin, Thielallee 63, 14195 Berlin, Germany; Randall Divison of Cell and Molecular Biophysics, King’s College London, London SE1 1UL, United Kingdom; Institut für Biochemie, ETH Zürich, Switzerland

## Abstract

The axon initial segment (AIS) is enriched in specific adaptor, cytoskeletal and transmembrane molecules. During AIS establishment, a membrane diffusion barrier is formed between the axon and the somatodendritic domain. Recently, an axonal periodic pattern of actin, spectrin and ankyrin forming 190 nm distanced, ring-like structures has been discovered. However, whether this structure is related to the diffusion barrier function is unclear.

Here, we performed single particle tracking timecourse experiments on hippocampal neurons during AIS development. We analyzed the mobility of lipid-anchored molecules by high-speed single particle tracking and correlated positions of membrane molecules with the nanoscopic organization of the AIS cytoskeleton.

We observe a strong reduction in mobility early in AIS development. Membrane protein motion in the AIS plasma membrane is confined to a repetitive pattern of ~190 nm spaced segments along the AIS axis as early as DIV4 and this pattern alternates with actin rings. Our data provide a new model for the mechanism of the AIS diffusion barrier.

## Introduction

Neurons are highly polarized cells that bear a complex dendritic arbor and usually only a single, elongated axon emanating from the cell body. The somatodendritic domain receives input from thousands of synapses and if a threshold is met, an action potential is generated by ion channels, which show a large density in the first 50-150 µm of the axon, the so-called axon initial segment (AIS) (Rasband, 2010). The AIS contains a specific complement of molecules with ankyrin G (AnkG), ßIV-spectrin, neurofascin and voltage-gated ion channels that is assembled during neuronal development (Galiano et al., 2012). The motion of membrane molecules in the AIS decreases in correlation with the accumulation of AnkG, an AIS-specific cytoskeletal adaptor molecule, during neuronal development until a diffusion barrier is established that impedes motion of axonal membrane molecules into the somatodendritic domain and vice versa (Winckler et al., 1999; Nakada et al., 2003; Boiko et al., 2007). This barrier is believed to be a result of the anchoring of transmembrane molecules to the submembrane cytoskeleton meshwork (Nakada et al., 2003). These tethered proteins then act as obstacles to the lateral motion of membrane molecules, which as a result may be transiently confined within the compartments of the meshwork – a general concept known as the picket fence model (Fujiwara et al. 2002).

Recent investigations have found a periodic cytoskeletal meshwork in the AIS (Xu et al., 2012; Zhong et al., 2014; Leterrier et al., 2015; d’Este et al., 2015), while electron microscopy studies have demonstrated that a dense coat of ion-channels and adhesion molecules is assembled at the AIS membrane during development (Jones et al., 2014). However, whether the structure of the periodic cytoskeletal meshwork specifically provides the basis for the membrane diffusion barrier, as in the picket fence model, or whether it is more generally induced by molecular crowding by a high density of membrane molecules along the AIS remains unclear (Rasband, 2013).

Here, we perform high-density single particle tracking (SPT) of glycosylphosphatidylinositol (GPI)-anchored molecules in the AIS over a timecourse of neuronal development and find that membrane protein motion is reduced as early as day in vitro 5 (DIV5). Using high-speed SPT, we go on to showing that the motion of GPI-anchored molecules is confined to a repetitive array of small areas spaced at ~190 nm. When we fix neurons after SPT experiments and perform super-resolution microscopy of the actin cytoskeleton, we find that the confinement areas and the equidistant 190 nm spaced actin rings reported by Xu et al. are mutually exclusive.

Our results suggest a new mechanism for the establishment of a membrane diffusion barrier in the AIS and demonstrate that the motion of GPI-anchored molecules in the plasma membrane can be controlled by the organization of the submembrane cytoskeleton.

## Results and discussion

To determine at what timepoint during neuronal development the AIS diffusion barrier is established, we performed timecourse experiments of high-density SPT on primary hippocampal neurons. To do so, we transfected neurons at DIV2 with GPI-GFP and then performed SPT using Atto647N-labeled anti-GFP nanobodies for the uPAINT method (Giannone et al., 2010) on the presumed AIS and a segment of the distal axon at DIV3 (Figure 1A). As the neurons were plated on gridded coverslips and experiments were performed in the presence of Cy3B-labeled anti-neurofascin antibodies, which is an AIS marker, we could identify the same neuron and AIS in subsequent SPT experiments at DIV5, 7 and 10. After the last experiment, we fixed the cells and performed immunofluorescence staining against AnkG to confirm the location of the AIS. When we analyzed the hundreds of obtained trajectories of single anti-GFP nanobodies in terms of the instantaneous diffusion coefficient (*D*), we found a decrease in *D* of GPI-GFP between DIV3 and DIV10 in the AIS (Figure 1B). However, we did not observe a decrease in the median *D* in the distal axon (Figure 1C, D) of the same neuron. Strikingly, we observed a significant reduction of the median *D* of GPI-GFP between DIV3 and DIV5 (Figure 1D) in the AIS of all tested neurons (N = 4 neurons, n = 3390 trajectories for DIV3, n = 11290 for DIV5, p = 0.003, paired t-test of the median *D*). This reduction was spatially correlated with the appearance of neurofascin-staining (Figure 1B, Supplementary Figure 1).

**Figure 1.**
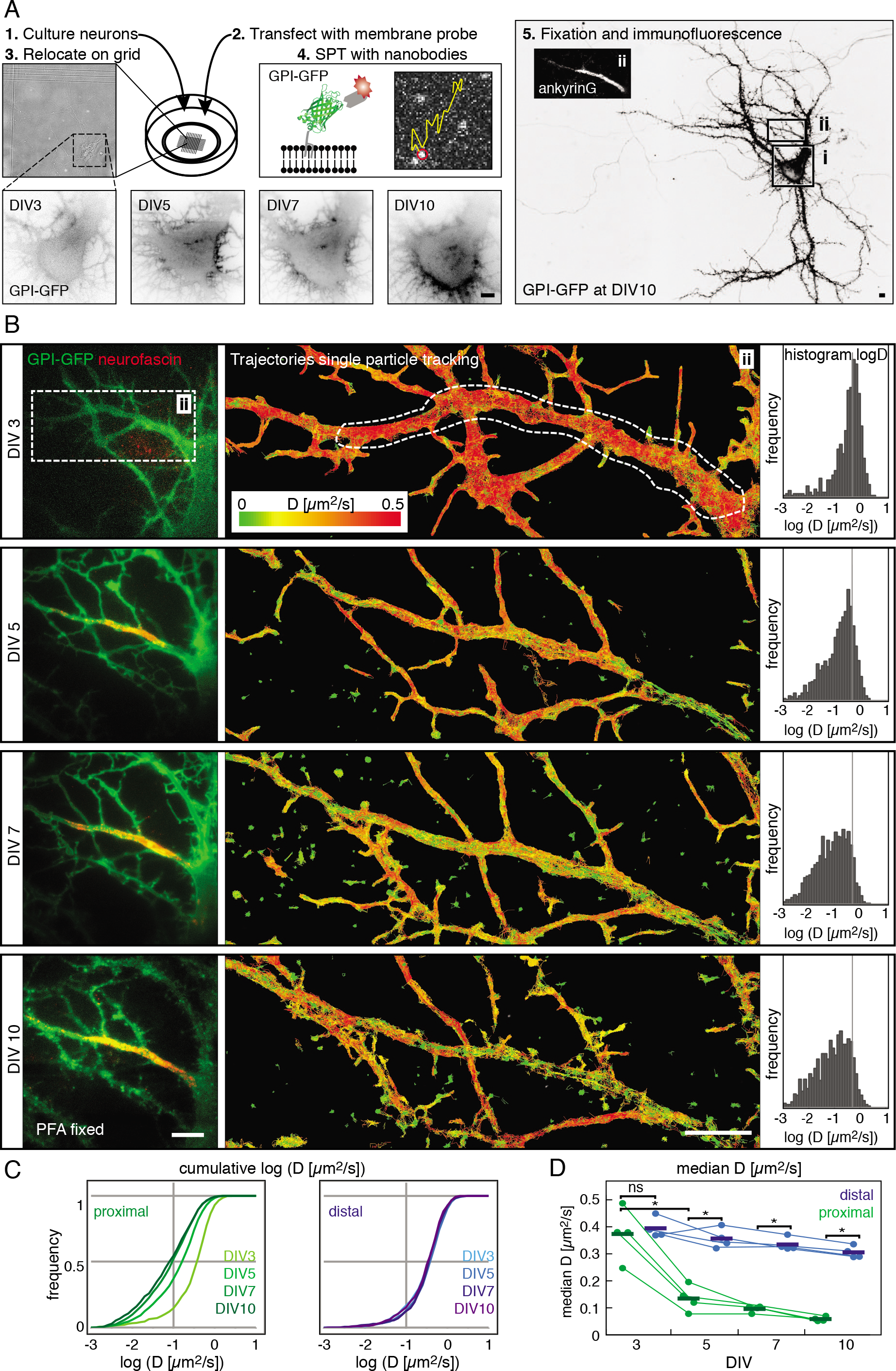
**The diffusion of membrane molecules is restricted by DIV5 in hippocampal neurons**. (A) Experimental design for the developmental timecourse in cultured primary hippocampal neurons. 1) Neurons were maintained in gridded glass-bottom petri dishes and 2) transfected with the membrane-probe GPI-GFP. 3) Individual neurons were located again on DIV3, 5, 7, and 10, respectively, based on their position on the grid. 4) Single particle tracking (SPT) experiments were conducted by sparsely labeling transfected neurons with anti-GFP nanobodies conjugated to fluorescent organic dyes. 5) Following SPT, neurons were fixed and immunostained for cytoskeletal and or AIS marker. The soma (i) and the proximal axon with the AIS (ii), identified by AnkG staining (inset micrograph), are outlined. (B) Results from the developmental SPT timecourse on a typical neuron. The AIS was identified by live-immunolabeling of neurofascin. Left box: The proximal axon was neurofascin negative on DIV3 and neurofascin positive from DIV5 onwards. Middle box: Plot of trajectories of mobile particles tracked in SPT experiments color-coded for instantaneous diffusion coefficients *D*. Right box: Histogram of *D* on the proximal axon where the AIS assembles (white dashed line). Number of trajectories: DIV3 n = 787, DIV5 n = 1815. DIV7 n = 2005, DIV10 n = 1067. (C) Plots of the cumulative *D* in the AIS (left, shades of green) and a portion of the distal axon (right, shades of blue) of the neuron shown in B. (D) Graph of the median *D* for all neurons (N = 4) between DIV3 and 10. Statistical analysis of median *D* by paired *t*-test, * p < 0.01. Scale bars: 5 µm.

Electron microscopy of the AIS of developing neurons does not show a significant accumulation of electron-dense material such as ion channels at the AIS at this early timepoint (Jones et al., 2014). On the other hand, the periodic submembrane spectrin-actin meshwork becomes detectable at this stage in neuronal development at the AIS (Zhong et al., 2014; d’Este et al., 2015) and individual ion channels start to become anchored by AnkG (Brachet et al., 2010). Our results thus suggested that the periodic submembrane spectrin-actin meshwork may be responsible for the observed reduction in diffusivity of GPI-GFP between DIV3 and DIV5 and not general crowding of the plasma membrane.

We therefore investigated the diffusivity of membrane molecules around DIV4 more closely as this seemed to be a transition point. We reasoned that if the periodic submembrane spectrin-actin meshwork was the cause of the decrease in diffusivity, a higher spatiotemporal resolution would be required in order to detect such a ~200 nm spaced periodic pattern in the diffusion of GPI-GFP. We estimated that motion blurring of molecules with high diffusivity such as GPI-GFP made it impossible to detect sharp compartmental borders with our single molecule detection-based method (Frost et al., 2012) and that SPT with millisecond temporal and nanometer spatial resolution was required to move forward. We thus decided to use streptavidin-coated quantum-dots (QDs), which are very bright and extremely photostable, and to couple them via biotinylated anti-GFP nanobodies to GPI-GFP for high-speed SPT. We reduced motion blurring by pulsing the excitation laser for 2 ms at 5-10 ms framerates on our EMCCD camera and used fiduciary markers to correct for drift (Figure 2A). By performing SPT measurements with a high density of membrane-bound QDs to achieve high sampling of the AIS membrane and recording tens of thousands of frames over several minutes, we generated trajectories of up to 2200 steps with a median localization accuracy of 8-15 nm (Figure 2B).

**Figure 2.**
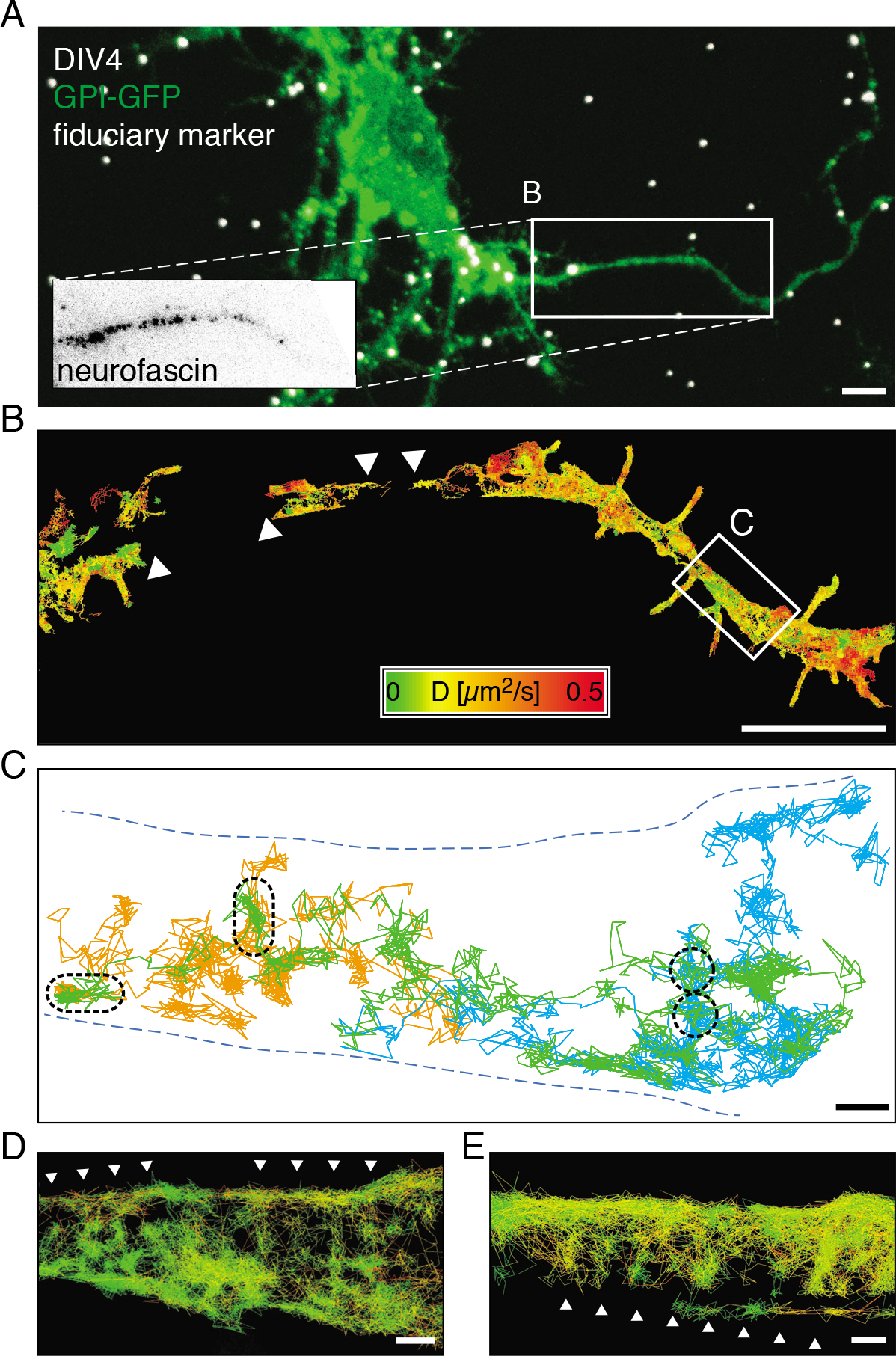
Highspeed single particle tracking shows that diffusing GPI-GFP molecules revisit small areas in the AIS. (A) Merged fluorescence micrograph of a DIV4 hippocampal neuron expressing GPI-GFP (green) post PFA fixation with fiduciary markers (white) for drift correction and image correlation. The AIS was identified by live immuno-labeling of neurofascin (box). (B) Plot of SPT trajectories of anti-GFP nanobody-coupled quantum dots (n = 3375) on GPI-GFP in the inset in A. Trajectories are color-coded according to the instantaneous diffusion coefficient. (C) Plot of selected long trajectories (> 500 frames, orange, green, blue) along a segment with reduced lateral mobility (white box in B). The trajectories cover the axon inhomogeneously and show local zones, which are revisited by individual QDs (black dashed lines). (D) Same plot of area as in C with all trajectories with more than 20 steps plotted (n = 231) and color-coded according to the instantaneous diffusion coefficient as in B. Arrowheads emphasize a pattern emerging from the distribution of the trajectories. (E) Plot of trajectories of anti-GFP nanobody coupled QDs on the AIS of a DIV11 neuron expressing GPI-GFP (n = 501). Trajectories are color coded as in B. Arrowheads emphasize an emergent pattern similar to that in D. Scale bars: 5 µm (A, B) and 200 nm (C, D, E).

We then analyzed subsets of the longest trajectories and found that specific areas were often revisited by individual QDs, suggesting that they preferred certain domains in the plasma membrane (Figure 2C). Indeed, when we plotted all measured trajectories, they seemed to form a striped pattern perpendicular to the direction of propagation of the axon both early in the developmental timecourse (DIV4, Figure 2D) and at later stages (DIV11, Figure 2E).

To analyze this pattern, we reconstructed images from the cumulative localizations of all detected QDs in all captured frames and found an alternating pattern perpendicular to the axon of zones with lower and higher localization densities (Figure 3A, C). When we calculated the spatial autocorrelation along the direction of propagation of the axon, we found a clear periodicity of ~200 nm, consistent with the spacing of the AIS cytoskeleton (Figure 3B, D).

**Figure 3.**
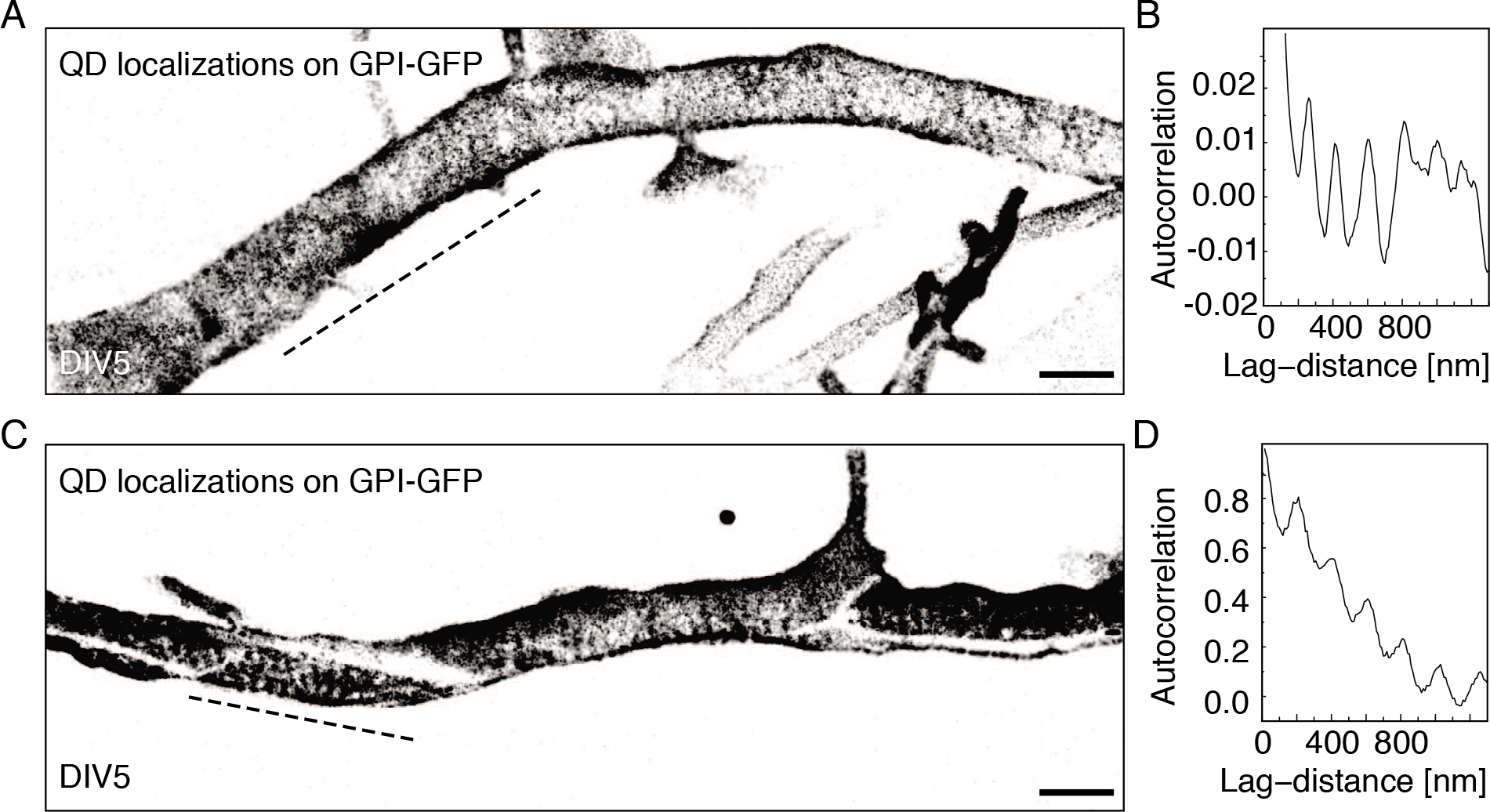
**Accumulated positions of GPI-GFP molecules from SPT show a periodic pattern of ~200 nm along the AIS**. (A) Reconstructed images from all localizations acquired during SPT of QDs on the proximal axon of a DIV5 hippocampal neuron expressing GPI-GFP. (B) Auto-correlation along the axon (dashed line) reveals a periodic pattern of ~200 nm. (C) The periodic pattern was observed along the entire proximal axon of some neurons but usually more prominent in some regions (dashed line). (D) Autocorrelation along a segment of axon (dashed line) with periodically arranged stripes of localizations. Scale bars: 500 nm.

The periodic submembrane spectrin-actin meshwork is established in the proximal axon as early as DIV2, before AIS-specific markers accumulate (Zhong et al., 2014; d’Este et al., 2015). Likewise, the earliest timepoint at which a periodic pattern emerged in an area showing low diffusion coefficient trajectories was DIV3 (Figure 4A-F). We performed many measurements during the developmental timecourse and on later timepoints and consistently found periodic patterns, albeit often only along segments that reached a sufficiently high localization density (Figure 4G-J, Supplementary Figure 2).

**Figure 4.**
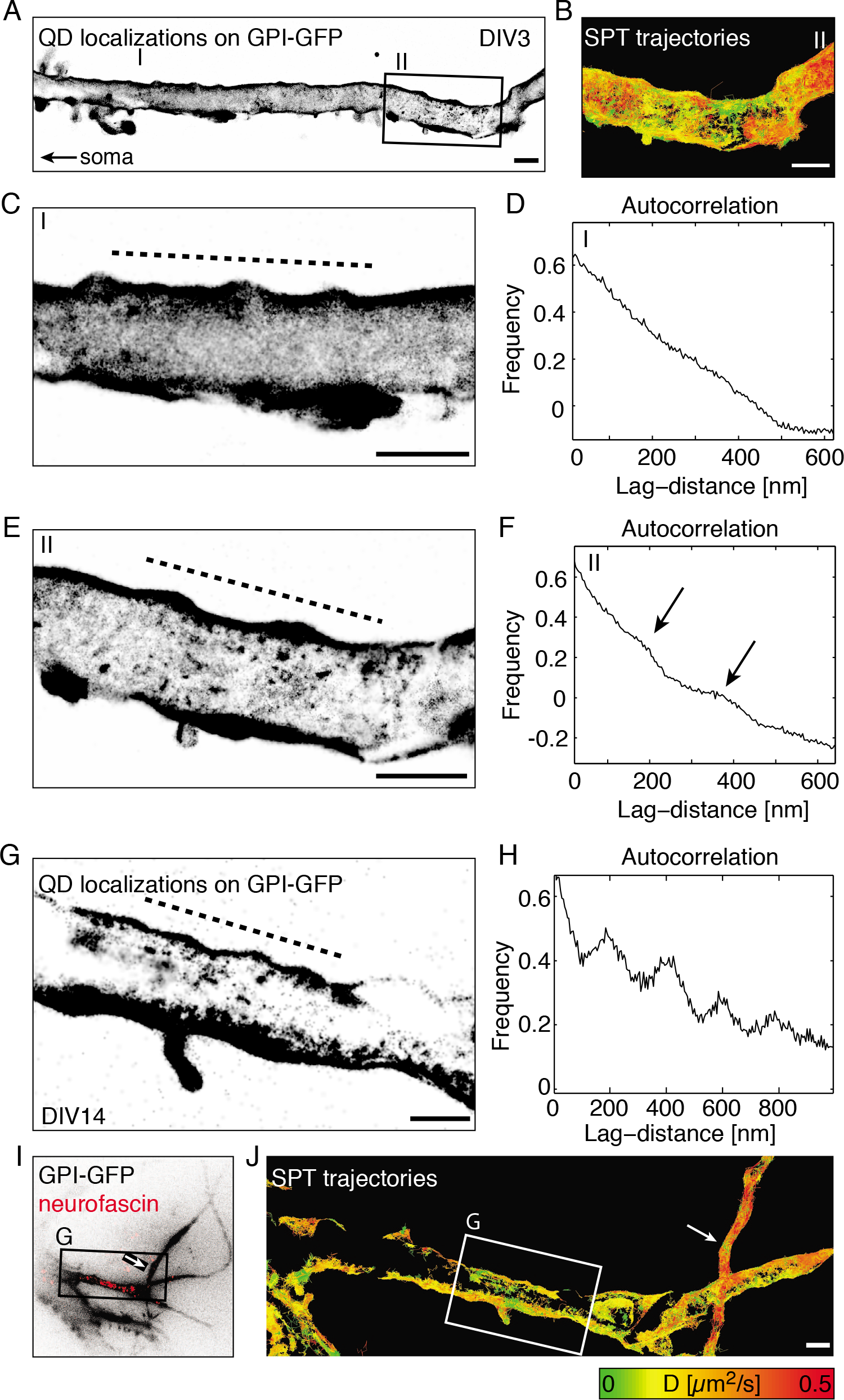
**The periodic confinement areas emerge as early as DIV3 during development**. (A) Reconstructed image of all localizations from SPT of QDs on GPI-GFP on a DIV3 hippocampal neuron show areas with randomly distributed localizations (I) adjacent to areas in which clusters of localizations form a pattern on the axon (II). (B) Trajectories color-coded for *D* show a reduced lateral mobility of QDs on GPI-GFP in regions with clustered localizations (II). (C) Close-up of (I) shows no visible pattern in localizations. (D) Auto-correlation along the axon (dashed line) shows no specific pattern in regions with homogeneously distributed localizations. (E) Close-up of (II) shows a segment of the axon with clusters of localizations. (F) Autocorrelation along this segment of the axon (dashed line) reveals an emerging periodicity of ~200 nm (arrows). (G) The periodic pattern was observed directly on the AIS in DIV14 neurons. (H) Auto-correlation of localizations (dashed line) clearly shows a periodicity of ~200 nm on the axon of this DIV14 neuron. (I) Overview of GPI-GFP signal with neurofascin live immunolabeling (red) that marks the AIS. (J) Trajectories color-coded for *D* on the neurofascin positive segment of the proximal axon show that the lateral mobility is reduced on the AIS in regions with periodic QD localizations compared to the distal axon that loops back to the neuron (arrow). Scale bars: 1 µm.

Our results strongly suggest that the global reduction of diffusion in the AIS was caused by an influence of the periodic submembrane spectrin-actin meshwork in the AIS. To investigate the spatial relationship between the periodic pattern of GPI-GFP localizations and the underlying cytoskeleton, we performed correlative SPT and super-resolution imaging of different components of the axonal cytoskeleton. For this, we fixed and immunostained neurons after SPT experiments to then perform super-resolution imaging of the cytoskeleton of the same AIS. First, we overlaid SPT measurements with dSTORM images of the C-terminus of ßII-spectrin, which is located at the center of the spectrin tetramer that connects the actin rings (Xu et al., 2012) (Supplementary Movie 1). We found extensive spatial overlap between the particle trajectories and ßII-spectrin immunostaining (Figure 5A-E). A spatial correlation analysis showed a strong cross-correlation between GPI-GFP and ßII-spectrin positions (Figure 5F).

**Figure 5.**
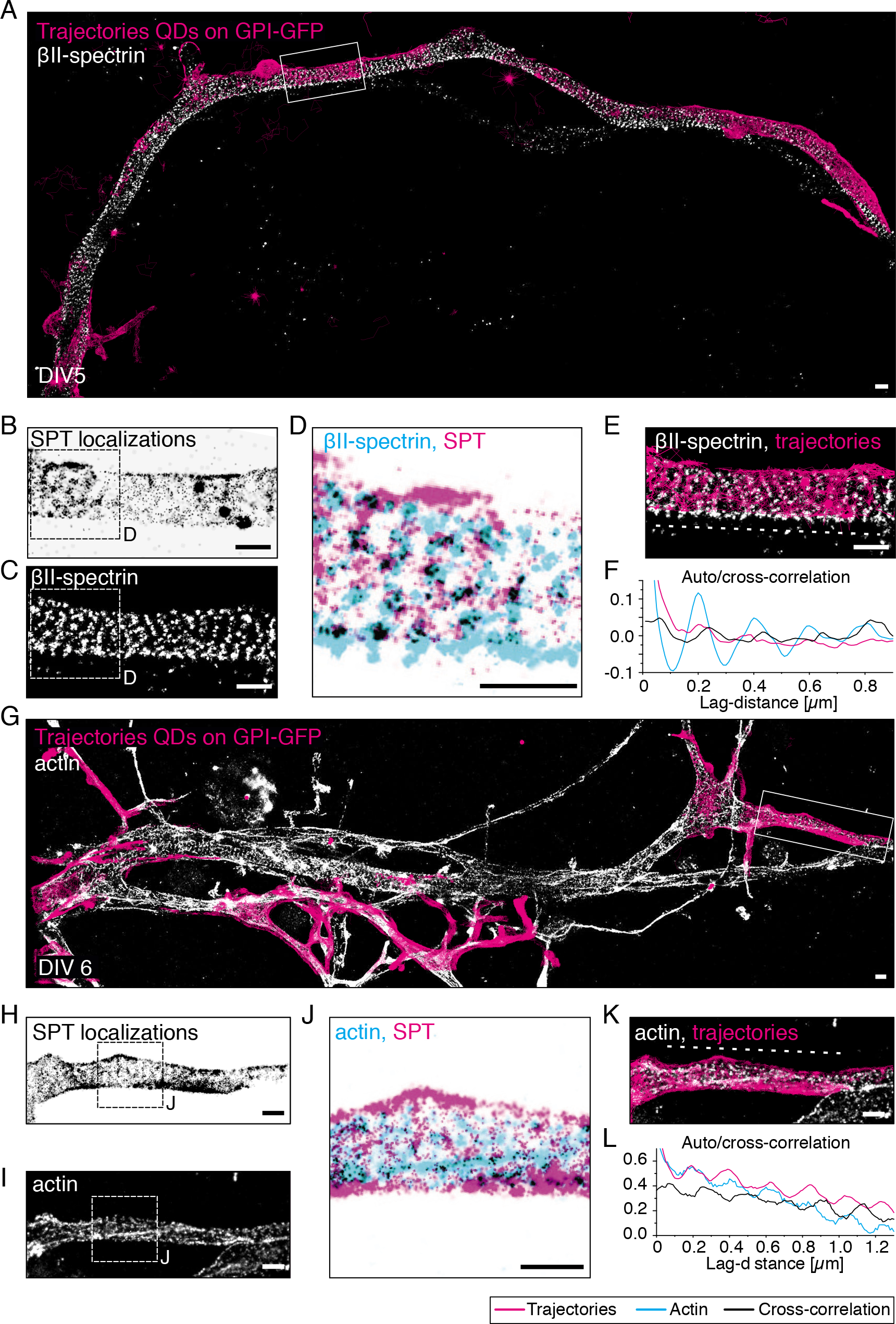
**The periodic confinement areas overlap with ßII-spectrin immunostaining and are excluded from actin rings**. (A) Proximal axon of a DIV5 hippocampal neuron with trajectories from SPT of GPI-GFP with QDs overlaid with a super-resolution micrograph of ßII-spectrin. (B) Reconstructed image of all localizations from SPT in region of interest (white box, A) shows a regular pattern along the axis of the axon. (C) Super-resolution micrograph of ßII-spectrin confirms periodicity of the submembrane cytoskeleton in the region of interest. (D) Zoom-in on region of interest (white box, B) with trajectories (magenta) and ßII-spectrin (white). (E) Overlay of localizations from SPT (magenta) and cytoskeletal marker ßII-spectrin (cyan). Regions where both overlap are black. (F) Auto- and cross-correlation along the axon (dashed line, E) confirms that both SPT localizations and ßII-spectrin are arranged periodically at ~200 nm. The cross-correlation places clusters of localizations from SPT on top of ßII-spectrin with an offset of ~25 nm. (G) Proximal axon of a DIV6 hippocampal neuron with trajectories from SPT of GPI-GFP with QDs overlaid with a super-resolution micrograph of actin. (H) Reconstructed image of all localizations from SPT in region of interest (white box, G) shows a regular pattern along the axis of the axon. (I) Super-resolution micrograph of actin confirms periodicity of the submembrane cytoskeleton in the region of interest. (J) Overlay of localizations from SPT (magenta) and cytoskeletal marker actin (cyan). Regions where both overlap are black. (K) Overlay of trajectories from SPT (magenta) and cytoskeletal marker actin (white). (L) Auto- and cross-correlation along the axon (dashed line, K) confirms that both SPT localizations and actin are arranged periodically at ~200 nm spacing. The cross-correlation places clusters of localizations from SPT in between the actin rings with an offset of ~20 nm. Scale bars: 500 nm.

When instead we correlated GPI-GFP trajectories with super-resolution images of actin using Alexa Fluor 647 phalloidin, GPI-GFP localizations and actin staining seemed mutually exclusive (Figure 5G-K, Supplementary Movie 2). Consistently, the cross-correlation analysis between GPI-GFP and actin showed a ~100 nm phase shift, corresponding to half a period of the cytoskeleton (Figure 5L). From this we concluded that GPI-GFP molecules are confined in their lateral motion in the neuronal plasma membrane between actin rings.

Taken together, our results show that between the third and the fifth day in culture of developing hippocampal neurons, a periodic array of physical barriers to the motion of GPI-anchored GFP in the plasma membrane is created in the AIS that localizes to actin rings. At this time, the full density of ion channels, adhesion molecules (Jones et al., 2014) and extracellular matrix (Frischknecht et al., 2009) at the AIS is not yet established, ruling out a major role of these assemblies in membrane protein motion. Earlier work has observed the establishment of a diffusion barrier in the AIS at DIV7-10 (Nakada et al., 2003; Boiko et al., 2007). However, the technological possibilities at the time allowed the observation of but few molecules.

Our work provides a mechanistic basis for the AIS diffusion barrier and raises a number of exciting questions regarding the organization and function of the neuronal cytoskeleton. It is known that the overexpression of ßII-spectrin induces a periodic cytoskeletal structure in dendrites (Zhong et al., 2014), and that dendrites stained with a fluorogenic analog of the actin-stabilizing drug Jasplakinolide exhibit periodic actin rings (d’Este et al., 2015). It will be important to investigate, whether such structures inhibit membrane protein motion as well and if the restriction of membrane protein motion in the AIS contributes to AIS maturation.

Beyond the development of the AIS, the finding that the motion of membrane molecules can be confined by the plasma membrane-associated cytoskeleton is important for all of cell biology. That the motion of a membrane molecule can be confined by actin corrals has long been proposed (Kusumi et al., 2005). Indeed it is clear that transmembrane molecules can be caught in a cytoskeletal meshwork (Tomishige and Kusumi, 1999) and that while lipids seem to undergo diffusion in the plasma membrane at the macroscopic scale, at the nanoscopic scale they exhibit subdiffusive motion (Fujiwara et al., 2002; Eggeling et al., 2009). In recent years, the organization of the plasma membrane and its control by cellular mechanisms is being investigated withsubmitted to new scrutiny using a number of novel experimental approaches. STED- FCS (stimulated emission depletion microscopy combined with fluorescence correlation spectroscopy) has demonstrated that the motion of lipids in the plasma membrane can be increased by an actin-depolymerizing drug (Andrade et al., 2015) and a combination of FCS with accurate temperature control of the sample showed that actin polymerization influences the motion of GPI-anchored proteins (Saha et al., 2015). However, these studies could not demonstrate a local correlation between actin and moving molecules and the mechanism behind these findings remains elusive. How a GPI-anchored molecule in the exoplasmic membrane leaflet of the plasma membrane can be confined by the submembrane cytoskeleton is subject of intensive investigation. It is known that transbilayer interactions of the alkyl chains of lipids chains can confine molecules in supported membrane bilayers (Spillane et al., 2014) and likely also in the plasma membrane of cells (Raghupathy et al., 2015). In the future, it would be exciting to investigate whether individual GPI-GFP molecules can be observed to be confined in protein-dense areas of the plasma membrane of other cultured cells as well (Saka et al., 2014).

Our work raises a number of questions regarding the function of the periodic axonal cytoskeleton and the role of the AIS diffusion barrier. In the future it will be exciting to investigate the functional consequences of membrane protein compartmentalization in the AIS on membrane transport and neuronal polarization and a possible compartmentalization of the dendritic and distal axonal plasma membrane.

## Materials and Methods

### Ethics statement

Humane killing for preparation of rat primary hippocampal neurons conformed to local King’s College London ethical approval under the UK Supplementary Code of Practice, The Humane Killing of Animals under Schedule 1 to the Animals (Scientific Procedures) Act 1986, or in accordance with the guidelines issued by the Swiss Federal Act on Animal Protection, respectively. All efforts were made to minimize animal suffering and to reduce the number of animals used.

### Cell culture

Primary hippocampal neurons were prepared from E18 Sprague-Dawley rats (Charles River) as previously described (Kaech and Banker, 2006). Neurons were maintained in neurobasal medium with B27 supplement and GlutaMAX (all Life Technologies) on poly-lysine (Sigma) coated 18 mm diameter #1.0 glass coverslips (Menzel) or µGrid glass bottom petri dishes (ZellKontakt GmbH) at 37 °C in a CO_2_-controlled humidified incubator. After three days the cytostatic cytosine β-D-arabinofuranoside (Sigma) was added at a final concentration of 5 µM. Neurons were transiently transfected with protein fusion constructs between DIV2 and DIV8 using Lipofectamine 2000 (Life Technologies). The GPI-GFP protein fusion construct was a kind gift from the Helenius laboratory (Keller et al., 2001).

### Single particle tracking

Anti-GFP nanobodies (Chromotek) were labeled with biotin-sulfo-NHS (Thermo Scientific) by standard N-hydroxysuccinimidyl ester chemistry according to the manufacturer’s protocol using a 3-fold molar excess resulting in approximately one biotin per nanobody. The labeled nanobody was purified from the excess of unreacted biotin using three 3 kDa MWCO desalting columns (Zeba Spin, Thermo Scientific). Biotin-anti-GFP nanobodies were conjugated to 1 nM Qdot 705 streptavidin (Thermo Scientific) at a ratio of 3:1 on the day of the experiment. Nanobody-conjugated QDs were added prior to image acquisition at a concentration of ~10 pM. A single image series of 25-50k frames, which took 2-5 min, was recorded with 2-10 ms exposure time and 2-3 ms pulsed laser illumination. The irradiance at the sample was ~2 kW/cm^2^. The EMCCD camera was run in its high-speed modality using the 20 MHz readout. 100 nm red-fluorescent (580/605) beads (Life Technologies) were added as fiduciary markers for drift correction and image correlation.

SPT using Atto647N-coupled anti-GFP nanobodies (Chromotek) was performed using the uPAINT method. Nanobodies were added immediately prior to image acquisition at a concentration of ~25 pM and multiple image series (typically 10-20) of 500 frames were recorded with 25 ms exposure time and 5 ms laser illumination time. A lower irradiance of ~0.5 kW/cm^2^ was used to reduce the extent of photobleaching.

SPT experiments were all carried out in live-cell imaging buffer (145 mM NaCl, 5 mM KCl, 10 mM Glucose, 10 mM HEPES, 2 mM CaCl_2_, 1 mM MgCl_2_, 0.2 % (w/v) BSA, 10 mM ascorbate (Ewers et al., 2014)) on a heated microscope stage.

The lateral localization accuracy was 25-30 nm for the tracking of Atto647N and 8-15 nm for the quantum dots. Instantaneous diffusion coefficients (*D_2-4_*) for individual trajectories were calculated from the mean square displacement (MSD).

### Immunofluorescence staining

Prior to fixation the neurons were briefly rinsed in warm PBS. For staining of ßII-spectrin and AnkG neurons were fixed for 15 min with 4 % (w/v) paraformaldehyde (Sigma) in PBS. For actin staining neurons were extracted and fixed with glutaraldehyde as previously described (Xu et al., 2012). Samples were permeabilized in 0.2 % (v/v) Triton-X100 (Sigma) for 30 min and blocked for 30 min with ImageIT (Life Technologies), then washed twice in PBS and subsequently blocked for 1 h in 10 % (v/v) horse serum, 1 % (w/v) BSA in PBS (blocking buffer).

As primary antibodies we used monoclonal mouse ßII-spectrin antibody (clone 42 Biosciences, targets a sequence close to the C-terminus of ßII-spectrin), monoclonal mouse AnkG antibody (clone 65, Neuromab) and monoclonal Neurofascin antibody (clone A12/18, Neuromab) conjugated in-house to Cy3 by NHS-ester chemistry. As secondary antibody we used donkey anti-mouse Alexa Fluor 647 (A31571, Life Technologies). Staining with antibodies was performed in blocking buffer using a 1 h incubation for primaries and secondaries, respectively. Actin filaments were labeled with Alexa Fluor 647 conjugated phalloidin (Invitrogen A22287) overnight at 4 ºC or ~1 h at room temperature. A concentration of ~0.5 µM phalloidin in blocking buffer was used.

### Microscope setups

Standard immunofluorescence microscopy was performed on a spinning-disk confocal microscope (inverted Olympus IX71, Roper Scientific). Super-resolution imaging and SPT experiments were performed on a custom-built setup. Briefly, a 473 nm laser (100 mW, Laserglow Technologies), a 556 nm (200 mW, Laserglow Technologies) and a 643 nm laser (150 mW, Toptica Photonics) were focused onto the back-focal plane of an Olympus NA 1.49, 60x, TIRF-objective. A quad-edge dichroic beamsplitter (405/488/532/635 nm, Semrock) was used to separate fluorescence emission from excitation light. Emission light was filtered by a quad-band bandpass filter (446/523/600/677 nm, Semrock) and focused by a 500 mm tube lens onto a back-illuminated EM-CCD chip (Evolve Delta, Photometrics).

### Single molecule localization microscopy

The buffer for super-resolution imaging consisted of 0.1 M MEA/0.2 M Tris, pH 8.0 with 5 % (*w*/*v*) glucose, 0.25 mg/ml glucose-oxidase and 20 µg/ml catalase. The imaging laser intensity of the 643 nm laser line used was ~2 kW/cm^2^. The intensity of the 473 nm activation laser was automatically adjusted to keep the average number of localizations per frame constant (maximum intensity ~0.5 kW/cm^2^). We recorded a minimum of 25’000 frames with an exposure time of 20-35 ms.

### Data analysis

Data analysis was performed in MATLAB (Mathworks). Positions of single particles were determined based on a maximum likelihood estimator by Gaussian fitting (Smith et al., 2010; Albrecht et al., 2015). Single molecule localization microscopy images were corrected for drift based on image-correlation while for SPT experiments the localizations were drift-corrected using fiducials. SPT data was registered with subsequent super-resolution images based on fiducials. We used a minimum of three fiducials and performed a rigid transformation based on the MATLAB built-in routine *cp2tform* using *non-reflective similarity*. We determined a fiducial registration error of under 25 nm.

## Acknowledgements

The authors acknowledge support from EU Marie Curie Action FP7-PEOPLE-2013-IEF 630024 (CMW) and Swiss National Fund Sinergia Grant CRS113_141945 (DA & HE).

## Author contributions

HE, CMW: designed research; DA, CMW: performed research; TT, CMW: contributed new reagents or analytic tools; CMW, DA, HE: analyzed data; HE: wrote the paper with input from all authors.

**The authors declare no conflict of interest**.

## Abbreviations

AIS: axon initial segment, AnkG: ankyrin G, *D*: instantaneous diffusion coefficient (*D_2-4_*), DIV: days in vitro, GPI-GFP: glycosylphosphatidylinositol-anchored green fluorescent protein, QD: quantum dot (semiconductor nanocrystal), SPT: single particle tracking.

## Supplementary Figures and Figure Legends

**Supplementary Figure 1.**
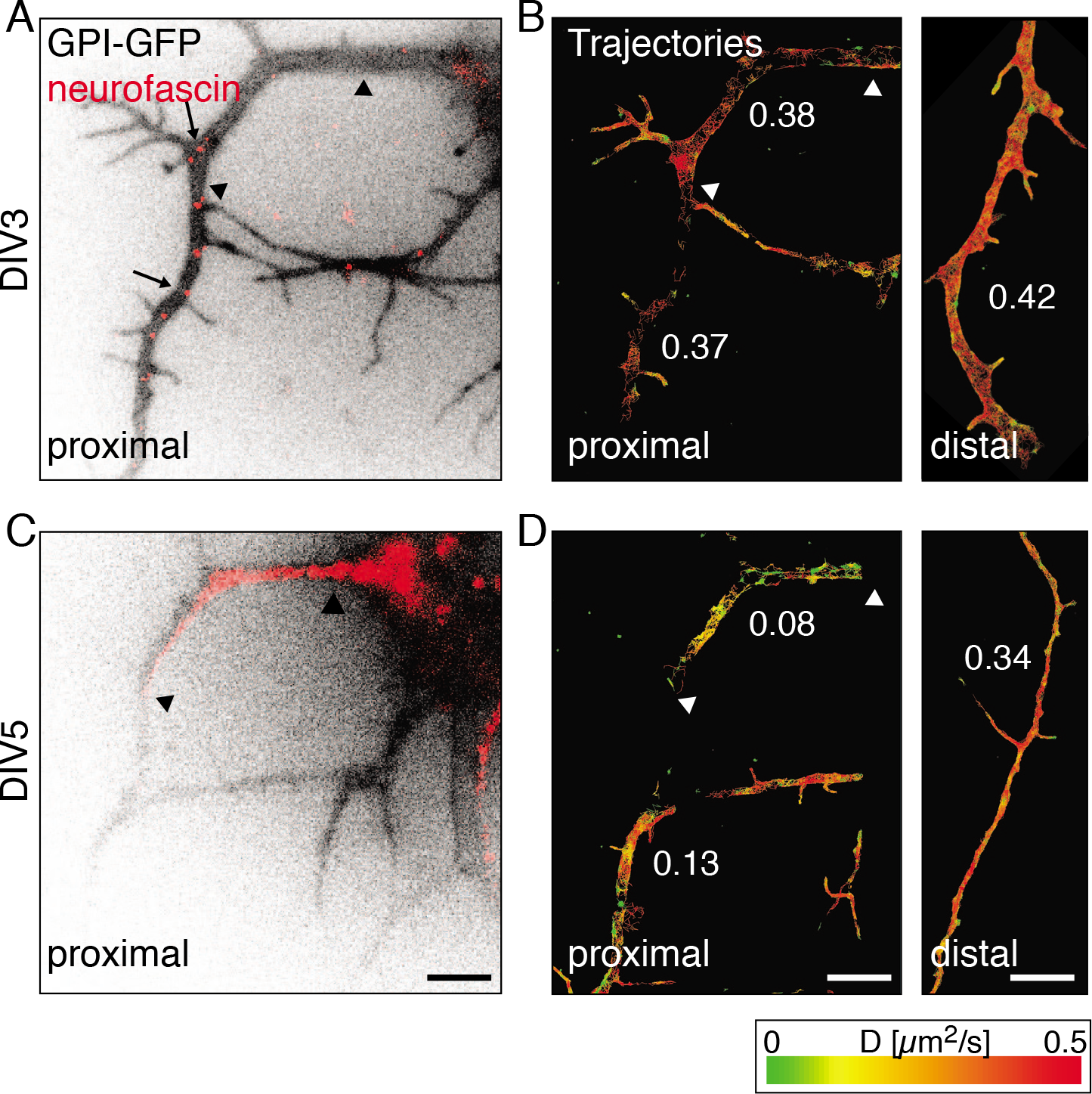
**The reduction of lateral mobility coincides with the accumulation of neurofascin at the AIS between DIV3 and DIV5**. (A) Shown is an area of the proximal axon of a DIV3 neuron expressing GPI-GFP. Live immunolabeling of neurofascin detected several neurofascin molecules along the proximal axon (arrows), however, not specifically localized at where the AIS later developed (arrowheads). (B) Trajectories from SPT of Atto647N-coupled anti-GFP nanobodies bound to GPI-GFP color-coded for D showed no difference along the proximal axon (0.38 vs 0.37 µm^2^/s, n = 451) on DIV3 and little difference compared to the distal axon (0.42 µm^2^/s, n = 1097). (C) On DIV5 the AIS was detectable by live neurofascin immunolabeling (arrowheads). (D) Trajectories color-coded for D showed that the lateral mobility was reduced in this region (0.08 µm^2^/s, n = 267) compared to the adjacent segment of the proximal axon without a clear neurofascin staining (0.13 µm^2^/s, n = 374). Particles in both regions exhibited a reduced lateral mobility compared to the distal axon (0.34 µm^2^/s, n = 1042) that was a little reduced compared to DIV3. Scale bars: 5 µm.

**Supplementary Figure 2.**
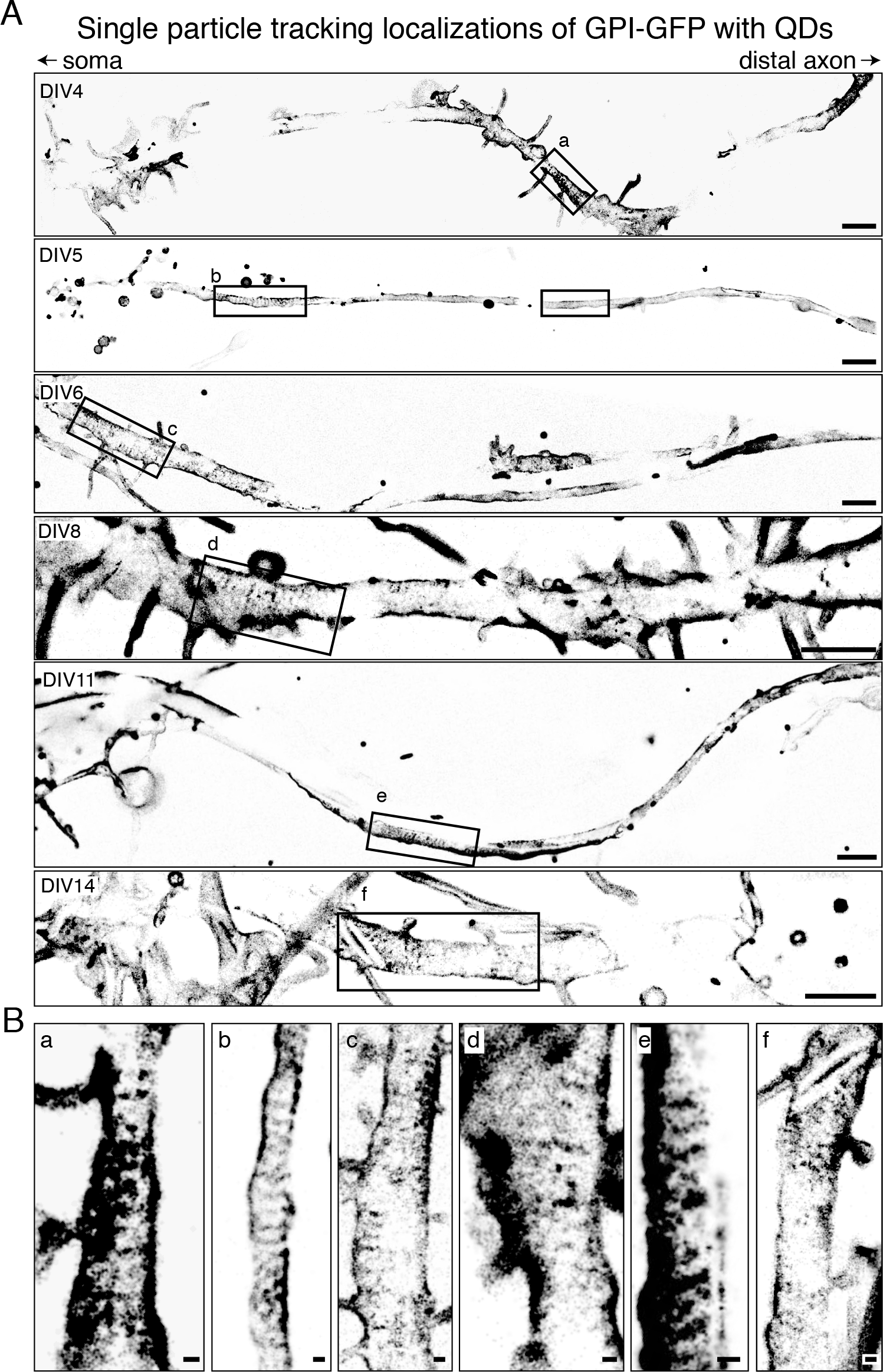
**Examples of the periodic pattern of localizations from SPT on the proximal axon over the timecourse of neuronal development**. (A) Neurons at DIV4 to DIV14 expressing GPI-GFP that was tracked with QDs on the proximal axon. Minimum 25’000 frames acquired. (B) Selected regions (a-f) with periodic stripes of localizations are enlarged. Notably, the pattern was not always observed continuously along the entire proximal axon but rather in segments with higher localization densities. Scale bars: 2 µm (A) and 200 nm (B).

**Supplementary Movie 1. ßII-spectrin super-resolution micrograph overlaid with trajectories of QDs on GPI-GFP in the axon initial segment at DIV5**. Shown is a movie assembled from 3652 frames with trajectories from connecting localizations in subsequent frames. Trajectories from SPT are projected in real time but sequentially and in random order on the super-resolution micrograph of ßII-spectrin. SPT data is correlated with the super-resolution micrograph based on fiduciary marker. Movie size is 2.25 by 7.5 µm. Time resolution is 12.1 ms. Playback speed is set to 83 frames per second (real-time).

**Supplementary Movie 2. Actin super-resolution micrograph overlaid with trajectories of QDs on GPI-GFP in the axon initial segment at DIV4**. Shown is a movie assembled from 7680 frames with trajectories from connecting localizations in subsequent frames. Trajectories from SPT are projected in real time but sequentially and in random order on the super-resolution micrograph of actin. SPT data is correlated with the super-resolution micrograph based on fiduciary marker. Movie size is 2.05 by 5.9 µm. Time resolution is 5.4 ms. Playback speed is set to 192 frames per second (real-time).

